# Evaluation of whole genome sequencing for the identification and typing of *Vibrio cholerae*

**DOI:** 10.1101/327973

**Authors:** David R. Greig, Ulf Schafer, Sophie Octavia, Ebony Hunter, Marie A. Chattaway, Timothy J. Dallman, Claire Jenkins

## Abstract

Epidemiological and microbiological data on *Vibrio cholerae* isolated between 2004 and 2017 (n=836) and held in the Public Health England culture archive were reviewed. The traditional biochemical species identification and serological typing results were compared with the genome derived species identification and serotype for a sub-set of isolates (n=152). Of the 836 isolates, 750 (89.7%) were from faecal specimens, 206 (24.6%) belonged to serogroup O1 and seven (0.8%) were serogroup O139, and 792 (94.7%) isolates from patients reporting recent travel abroad, most commonly to India (n=209) and Pakistan (n=104). Of the 152 isolates of *V. cholerae* speciated by kmer identification, 149 (98.1%) were concordant with the traditional biochemical approach. Traditional serotyping results were 100% concordant with the whole genome sequencing (WGS) analysis for identification of serogroups O1 and O139 and Classical and El Tor biotypes. *ctxA* was detected in all isolates of *V. cholerae* O1 El Tor and O139 belonging to sequence type (ST) 69, and in *V. cholerae* O1 Classical variants belonging to ST73. A phylogeny of isolates belonging to ST69 from UK travellers clustered geographically, with isolates from India and Pakistan located on separate branches. Moving forward, WGS data from UK travellers will contribute to global surveillance programs, and the monitoring of emerging threats to public health and the global dissemination of pathogenic lineages. At the national level, these WGS data will inform the timely reinforcement of direct public health messaging to travellers and mitigate the impact of imported infections and the associated risks to public health.

## Introduction

Cholera is an acute diarrhoeal disease that can kill within hours if left untreated. Patients present with the passing of voluminous rice water stools leading to severe dehydration (1). If hydration and electrolyte therapy is not quickly initiated, symptoms can rapidly progress to hypovolemic shock, acidosis and death. Inadequate access to clean water and sanitation facilities is a driver of transmission, and outbreaks are common among displaced populations living in overcrowded conditions (2).

The bacterial pathogen responsible for the disease is *Vibrio cholerae. V. cholerae* serogroups O1 and O139 are regarded as pandemic strains and harbour the *ctx* genes associated with the production of cholera toxin (3). *ctx* has also been detected in a limited number of other serogroups (4). Serogroup O1 can be divided into two biotypes, Classical and El Tor. There are over 200 different lipopolysaccaride ‘O’ antigens or serogroups of *V. cholerae*. The non-O1, non-O139 serogroups are associated with a milder form of gastroenteritis, septicaemia and other extra-intestinal infections (1, 3).

Seven cholera pandemics have occurred throughout the 19th and 20^th^ centuries. The seventh (and current) pandemic began in the Bay of Bengal and has spread to Africa and South America in at least three independent but overlapping waves of transmission (5). The fifth and sixth pandemics were caused by the *V. cholerae* serogroup O1 Classical biotype while the seventh pandemic was caused by serogroup O1 biotype El Tor. In 1992, *V. cholerae* serogroup O139 caused a large epidemic in Bangladesh and India (6), however *V. cholerae* O1 El Tor persists as the most commonly isolated ctx-positive serotype/biotype. *V. cholerae* is endemic across Africa, Latin America and Asia resulting in a large healthcare burden in developing countries (7–9). The World Health Organisation states that there are 1.3 - 4 Million estimated cases and 21,000- 147,000 estimated deaths annually (7).

The UK Standards for Microbiology Investigations Investigation of Faecal Specimens for Enteric Pathogens recommends testing of faecal specimens for *V. cholerae* in cases of suspected cholera, seafood consumption, and/or recent travel (within 2-3 weeks) to countries where cholera is endemic (https://www.gov.uk/government/publications/smi-b-30-investigation-of-faecal-specimens-for-enteric-pathogens). Consequently, the true incidence of domestically acquired *V. cholerae* in the UK is unknown, and almost all isolates of enteric origin are associated with travellers’ diarrhoea.

In 2015, Public Health England (PHE) implemented whole genome sequencing (WGS) for the routine surveillance of the more common gastrointestinal pathogens including *E.coli, Salmonella, Campylobacter, Shigella* and *Listeria* species (10–12). The aim of the study was to review the historical PHE data on isolates of *V. cholerae* held in the PHE culture archives, compare the results of the traditional biochemical and serological methods with the analysis of WGS data for a sub-set of isolates, and assess the impact of implementing WGS for the public health surveillance of *V. cholerae*.

## Methods

### Epidemiological data

All isolates of *V. cholerae* from human cases resident in England submitted to the Gastrointestinal Bacteria Reference Unit (GBRU) by local hospital laboratories between 2004 and 2017 were reviewed. Patient information including, sex, age and recent travel, was collected from laboratory request forms upon submission and stored in the Gastro Data Warehouse (GDW), an in-house PHE database for storing and linking patient demographic and microbiological typing data. Data on symptoms were limited stating only that the patient had either gastrointestinal symptoms or an extra-intestinal infection. There was no data on severity of symptoms or patient outcome.

### Bacterial culture and traditional biochemistry and serology

Cultures were stored on cryobeads at −40°C or in nutrient agar stabs. For each sample, one cryobead was taken and inoculated into 20ml 3% NaCl peptone water and incubated at 37°C for 18 hours, shaking at 80rpm. Cultures were plated out from either nutrient agar slopes or 3% NaCl peptone water onto Blood agar, MacConkey agar with salt (NaCl 1%), Thiosulphate-citrate-bile salts (TCBS) agar and cystine lactose electrolyte deficient (CLED) agar and incubated at 37°C for 18 hours.

Biochemical identification was performed following inoculation onto a panel of substrates. Utilisation of the substrate was identified by a colour changes or gas production within the media. All positive and negative reactions were compared to a known reference panel of results to give a final identification. Isolates of *V. cholerae* were agglutinated with antisera raised to O1 (Ogawa and Inaba) and O139 (Bengal) antisera to determine the serogroup. The Classical and El-Tor biotypes were differentiated by the Voges-Proskauer (VP) test (Classical, negative; El-Tor, positive) and haemolysis on blood agar (Classical, non-haemolytic; El-Tor, haemolytic).

### Whole genome sequencing analysis

All viable cultures of *V. cholerae* submitted to GBRU between January 2015 and March 2018 were sequenced (n=152). Genomic DNA was extracted, fragmented and tagged for multiplexing with Nextera XT DNA Sample Preparation Kits (Illumina) and sequenced using the Illumina HiSeq 2500 at PHE. FASTQ reads were quality trimmed using Trimmomatic (v0.36) (13) with bases removed from the trailing end that fell below a PHRED score of 30. If the read length post trimming was less than 50, the read and its pair were discarded using Trimmomatic. FASTQ reads from all sequences in this study can be found at the PHE Pathogens BioProject at the National Center for Biotechnology Information (PRJNA438219).

A kmer (a short string of DNA of length k; in this method k=18) based approach was used to confirm the identity of the sample before organism specific algorithms were applied (https://github.com/phe-bioinformatics/kmerid) (14). Reference genomes (n=1781) in 59 bacterial genera comprising the majority of human pathogens, commensal bacteria and common contaminants were downloaded from ftp://ftp.ncbi.nlm.nih.gov/genomes/refseq/bacteria. The kmer algorithm compared each sample to representative genomes in these 59 bacterial genera and returned the most similar genome together with a similarity estimate.

Sequence Type (ST) assignment was performed using a modified version of SRST using the MLST database described by Tewolde *et al.* 2016 (15). The MOST software (for MLST) is available at https://github.com/phe-bioinformatics/MOST. Any MLST gene sequences that did not match the existing alleles were submitted to pubMLST (https://pubmlst.org/vcholerae/) for a new allelic type assignment. Similarly, new allelic profiles were also submitted to the database for a new sequence type (ST) assignment.

For the isolates belonging to clonal complex (CC) 69, high quality Illumina reads were mapped to a SPADES v3.5.0 *de novo* assembly of the *V. cholerae* reference genomes NC-002505.1 and NC-002506.1, using BWA-MEM v0.7.3 and Samtools v1.1 (16, 17). Single Nucleotide Polymorphisms (SNPs) were identified using GATK v2.6.5 (18) in unified genotyper mode. Core genome positions, defined as those present in the reference genome and at least 80% of the isolates, that had a high quality SNP (>90% consensus, minimum depth 10x, MQ >= 30) in at least one isolate were extracted using SnapperDB v0.2.5 and processed though Gubbins v2.0.0 to account and supress recombination within the input to RAxML v8.1.17 (19).

Using the *GeneFinder* tool (Doumith, unpublished), FASTQ reads were mapped to the virulence regulator gene, *toxR* (Genbank accession: KF498634.1), the cholera toxin gene *ctxA* (Genbank accession: AF463401.1), *wbeO1*, and *wbfO139* (Genbank accessions: KC152957.1 and AB012956.1) encoding the somatic O antigens O1 and O139, *tcpA* classical and *tcpA* El Tor gene sequences (Genbank accessions: M33514.1 and KP187623.1) using Bowtie 2 (20). The best match to each target was reported with metrics including coverage, depth and nucleotide similarity in XML format for quality assessment. *toxR* is found in all isolates of *V. cholerae* and are regarded as a marker for species identification, *ctxA* encoding cholera toxin is associated with *V. cholerae* O1 and O139 and is a marker for the pandemic lineages (21). Variants of *tcpA* can be used to identify the Classical and El Tor biotypes (21). For *in silico* predictions, only results that matched to a gene determinant at >80% nucleotide identity over >80% target gene length were accepted.

## Results

### Review of the historical data

Between January 2014 and December 2017, 836 isolates of *V. cholerae* from human cases resident in England were submitted to GBRU by local hospital laboratories. On average, the number of isolates per year was 60, with the lowest number of isolates being reported in 2013 (n=29) and the highest number was reported in 2007 (n=80) (Figure 1). Of the 836 isolates, 206 (24.6%) belonged to serogroup O1 and seven were serogroup O139 (0.8%), and 750 were from faecal specimens, six were from blood cultures, two were from ear swabs and two were from eye swabs. No clinical data was available for the remaining 76 isolates.

**Figure 1.**
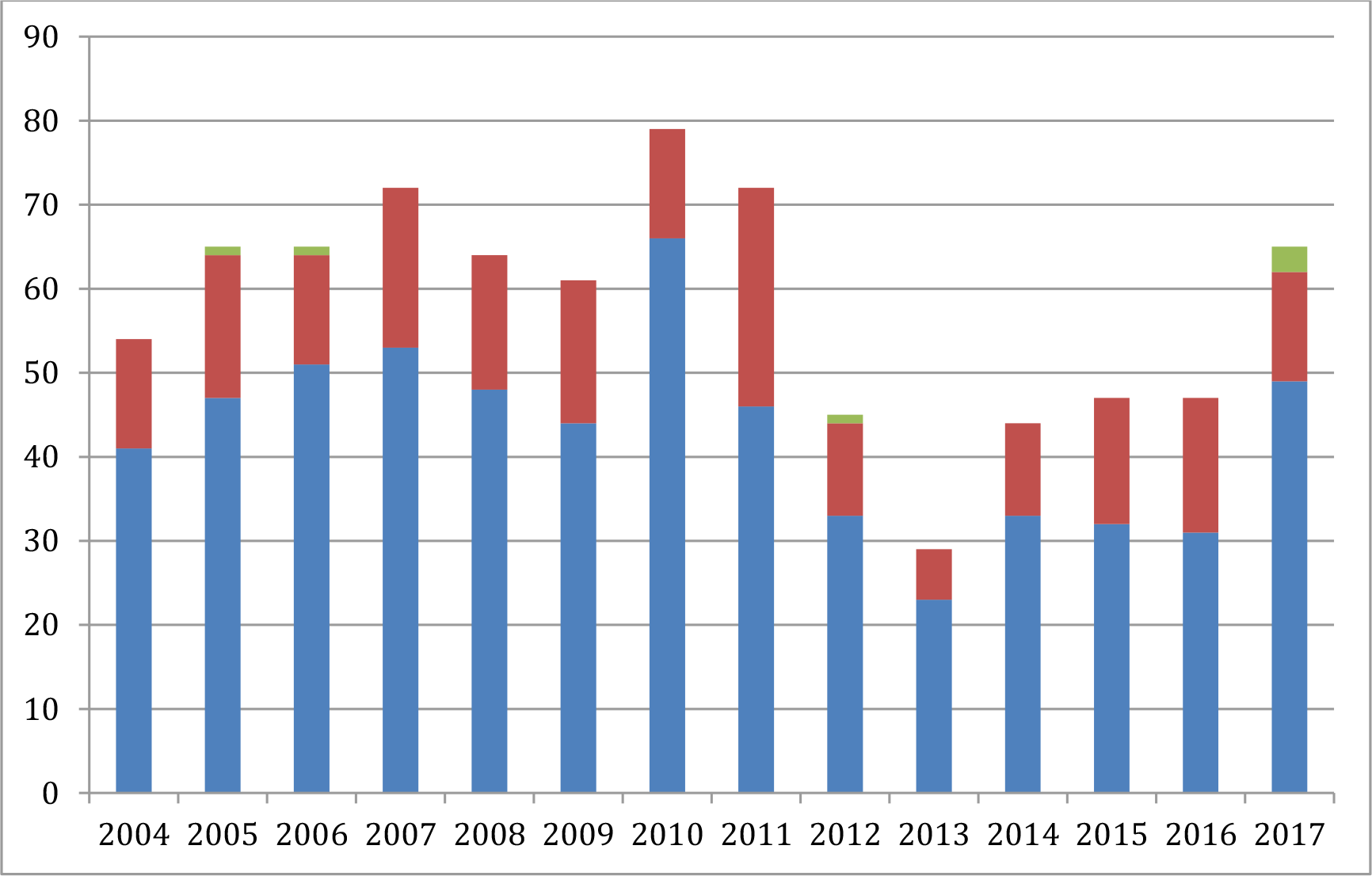
Number of isolates of *V. cholerae* from human cases resident in England submitted to GBRU by local hospital laboratories each year between 2004 and 2017 (n=836). Non-O1, non-O139 serogroups - blue; Serogroup O1 - red; Serogroup O139 - green

**Figure 2.**
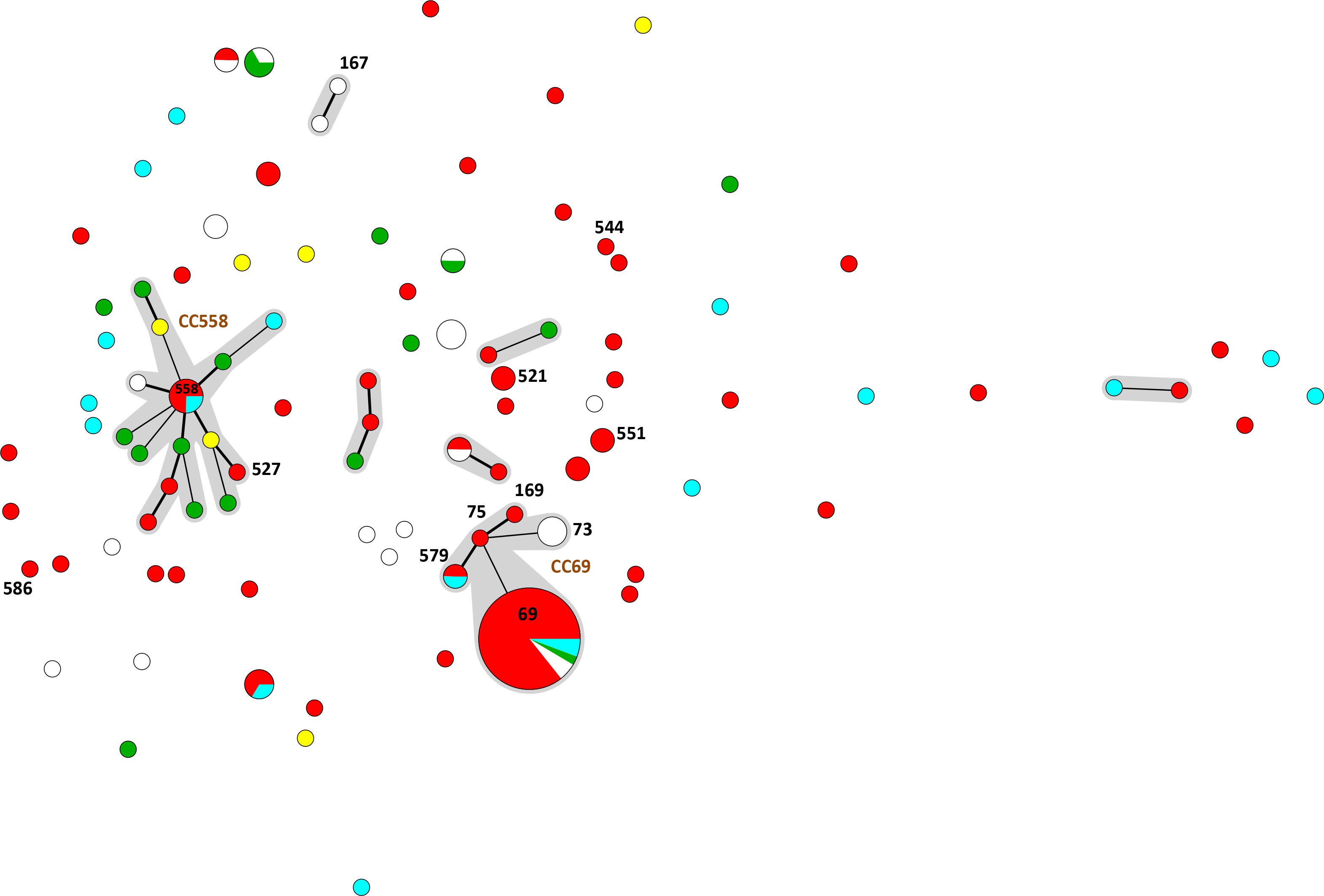
Minimum spanning tree illustrating the diversity in the population structure of the isolates of *V. cholerae* received at PHE between 2015 and 2017. Clonal complexes (CC) comprising strains linked by a single locus variant (thick black line) or double locus variant (thin black line) and are shaded grey. Sequence types (ST) are shown in black. Isolates associated with cases reporting recent travel abroad are highlighted: red - Asia; blue - Africa, green - Latin America, yellow - mainland Europe, while - no data.

Gender and age data was available for 828/836 and 773/836 cases, respectively. There were 424/836 males (50.7%) and 404/836 females (48.3%), and 685/836 (81.9%) were adults (aged 16 years or older) and 88 (10.5%) were children (<16 years old). Travel data was available for 796/836 cases, of which 792 reported recent travel abroad (less than 7 days of onset of symptoms). For the cases infected with *V. cholerae* non-O1, non-O139, the most common travel destinations were India (n=140), Kenya (n=57), Thailand (n=40) and Egypt (n=36). The most commonly reported destinations of cases of *V. cholerae* O1 were Pakistan (n=72) and India (n=69). The six cases of *V. cholerae* O139 had travelled to Thailand (n=2), China, India, Jordan and Pakistan. The four cases who stated they had not recently travelled abroad, all had *V. cholerae* non-O1, non-O139 isolates from extra-intestinal sites (blood cultures, n=2; eye swab, n=1; ear swab, n=1).

### Whole genome sequencing

One hundred and fifty-two isolates of *V. cholerae* were sequenced including those belonging to serogroup O1 (n=47), serogroup O139 (n=7), those designated serogroup non-O1, non-O139 (n=98) (Supplementary Table 1). One hundred and thirty-seven were isolates from human cases, of which 132 were from faecal specimens from hospitalised or community cases with symptoms of gastrointestinal disease, three isolates were from ear swabs, one was from an eye swab and one was from a blood culture from a patient with acute cholecystitis. The remaining 15 isolates were from animals (n=4), food (n=1), environmental samples (n=3) or were isolates from the National Collection of Type Cultures (n=7) (Supplementary Table 1).

### Kmer identification

Of the 152 isolates of *V. cholerae* speciated using the kmer identification approach, 149 (98.1%) were concordant with the traditional biochemical identification. The kmer method failed to identify three unusual external quality assessment isolates from an obscure environmental source, previously identified as *V. cholerae.* These isolates were *V. cholerae*, however, the similarity of the sequences to the *V. cholerae* reference sequences in the kmer ID database was below the acceptable threshold (80% similarity) required to confirm the identification.

### Use of Genefinder for serotyping and detection of virulence genes

The *toxR* gene was detected in 144/152 isolates of *V. cholerae* (Table 1). Eight isolates identified as *V. cholerae* by traditional biochemical tests, were negative for *toxR.* Further analysis of the sequences data showed the sequence coverage/similarity of *toxR* for the discrepant isolates fell just below the 80% threshold (74% and 77%) (Table 1). However, all eight isolates were identified at *V. cholerae* by the kmer approach.

**Table 1.** Summary of GeneFinder profiles and ST results

| Genefinder profile | ST | Number of isolates |
| --- | --- | --- |
| *toxR, wbeO1, tcp Classical, ctxA* | 73 | 3 |
| *toxR, wbeO1, tcp El Tor, ctxA* | 69 | 34 |
| *toxR, wbeO1, tcp El Tor* | 75, 169, 579 (2) | 4 |
| *toxR, wbeO1, ctxA* | 167 | 1 |
| *toxR, wbeO1* | 521(2), 551 (2) | 4 |
| *toxR, wbfO139, tcp El Tor, ctxA* | 69 | 1 |
| *toxR, wbfO139* | 163, 527, 529, 544, 568 | 5 |
| *toxR (77%), wbfO139* | 586 | 1 |
| *toxR* | >70 different STs | 94 |
| *toxR (77%)* | 539, 540, 541, 550, 585, 587, 600 | 7 |
|  | Total | 152 |

Traditional biochemistry and serotyping results were 100% concordant with the WGS analysis for identification of O1 and O139 and Classical and El Tor biotypes. Of the 37 isolates of *V. cholerae* O1 El Tor, 33 had *wbeO1, tcpA* El Tor variant and *ctxA* and four had had wbeO1, *tcpA* El Tor variant without *ctxA* (Table 1). There were five isolates that belonged to *V. cholerae* serogroup O1 that were negative for *tcpA* El Tor variant and *ctxA*, and one that was negative for *tcpA* El Tor variant but had *ctxA* (Table 1). There were only three Classical variant strains in the study, as the current pandemic is caused by *V. cholerae* O1 El Tor (7–9, 22, 23). All three *V. cholerae* O1 classical strains had *wbe*O1 and the *tcpA* classical variant gene. *V. cholerae* O139 was also rare in this dataset (24). Although all seven isolates of *V. cholerae* O139 had *wbf*O139, only the NCTC strain had the *tcpA* El Tor variant and *ctxA* (Table 1).

*ctxA* was detected in all isolates of *V. cholerae* O1 El Tor and O139 belonging to ST69 (25) and *V. cholerae* O1 Classical variants belonging to ST73 (22). Four isolates of *V. cholerae* O1 were negative for *ctxA*, and the six recently isolated of *V. cholerae* O139, were *ctxA*-negative.

### Sequence typing

Sequence typing data was available for 152 isolates. The *V. cholerae* O139 El Tor -positive isolate and the 34 isolates of *V. cholerae* O1 El Tor *ctxA*-positive isolates belonged to ST69. The four *V. cholerae* O1 El Tor *ctxA*-negative isolates belonged to ST75, ST169 and ST579 (n=2) and all fell within CC69. Previously studies have suggested that the emergence and potential spread of ST75 may pose significant threat to public health (26). Epidemiological surveillance is required to further understand the epidemic potential of *ctxA*-negative STs that are part of CC69. The *V. cholerae* O1 classical isolate belonged to ST73.

The six isolates of *V. cholerae* O1 without the *tcpA* El Tor variant were ST167, ST521 (n=2) and ST551 (n=2) and ST611. There were six isolates that had the O139 antigen but were negative for the *tcpA* El Tor variant gene and *ctxA*, and these belonged to ST163, ST527, ST529, ST544, ST568 and ST586. All *V. cholerae* O1 isolates, regardless of the presence or absence of *tcpA* El Tor variant or *ctxA*, belonged to the CC69 cluster and those without the tcpA El Tor variant gene were dispersed across the population (Figure 1).

The remaining 95 isolates of *V. cholerae* non-O1, non-O139 (n=95) and *V. cholerae* O139 (n=3), belonged to over 70 different STs (Supplementary Table). There was only one major cluster among the *V. cholerae* non-O1, non-O139 isolates, designated CC558. Isolates belonging to this cluster were geographically dispersed (Figure 1).

### SNP typing

As previously described, the pandemic *V. cholerae* O1 and O139 El Tor *ctxA* strains all belonged to ST69, whereas the Classical biotype and *ctxA*-negative strains of *V. cholerae* O1 belonged to other STs within CC69. A phylogeny of ST69, the pandemic lineage, was constructed comprising isolates from this study and sequences available in public databases (Supplementary Figure). The isolates from UK travellers clustered geographically with those returning from India located on the same branch, and those reporting recent travel to Pakistan clustered on a separate branch. Further analysis based on single nucleotide polymorphisms in the core genome compared to a reference strain may be performed for outbreak detection and source attribution where the incidence of the current *V. cholerae* O1 El Tor pandemic lineage (ST69) is high (27, 28)

## Discussion

Historically, traditional biochemistry, biotyping, phage typing and serology results were useful for confirming identification at the species level, typing of serogroups O1 and O139 and for identifying the Classical and El Tor variants. Isolates belonging to serogroups O1 and O139 were assumed to belong to the pandemic lineages and have the potential to cause cholera. In this study, the review of the historical GBRU data revealed that just under a quarter of the isolates of *V. cholerae* belonged to serogroup O1 and *V. cholerae* O139 was rarely detected (22). Due to limited resources, neither serotyping of the non-O1, non-O139 serogroups, nor molecular typing of any serogroup, were performed at GBRU. Therefore, prior to the implementation of WGS, it was not possible to monitor trends in emerging pathogenic lineages or gain insight into modes of transmission for this important gastrointestinal pathogen.

Previous studies have shown that MLST data is an accurate, robust, reliable, high throughput typing method that is well suited to routine public health surveillance (11, 12, 15). For *V. cholerae*, MLST provides insight on the true evolutionary relationship between isolates, as well as a framework for fine level typing for public health surveillance (29–32). Using the STs derived from the genome data, we were able to analyse the population structure of all isolates of *V. cholerae* submitted to GBRU for the first time. As previous studies have shown, the population structure of the non-O1 and non-O139 serotypes was diverse (31). Currently, the correlation of ST with geography in our dataset is hindered by the limited size of the dataset. However, moving forward this unprecedented level of strain discrimination available for all isolates of *V. cholerae* submitted to GBRU will enhance our understanding of global transmission and emerging threats to public health, for the pandemic strains belonging to CC69, and the non-O1 serogroup lineages.

A review of the data on cases of travellers’ diarrhoea caused by *V. cholerae* held by GBRU showed that travel histories, including the country visited, were complete for 95.2% of cases. Therefore, these data have the potential to be a useful public health resource for global surveillance, enabling us to track the emergence and dissemination of specific lineages on a global scale (5, 8, 33). Furthermore, at the national level, sharing of WGS data linked to these cases could result in the timely reinforcement of direct public health messaging to travellers, in order to reduce the number of imported infections and mitigate the impact of imported infections and associated risks to public health (34).

The consequence of humanitarian crises, such as the disruption of water and sanitation systems and the displacement of populations to overcrowded camps, increases the risk of the transmission and outbreaks of cholera (2). However, robust global monitoring of *V. cholerae* is hindered by the limitations of the surveillance systems in countries where people are most at risk. The World Health Organisation recommends that cholera surveillance should be part of an integrated disease surveillance system that includes feedback at the local level and information sharing at the global level (http://www.who.int/mediacentre/factsheets/fs107/en/).

Traditional biochemistry and serotyping results were concordant with the WGS analysis for identification of *V. cholerae* O1, serotyping and biotyping of O1 and O139 serogroups. Moreover, using the WGS approach species level identification, serotyping, biotyping, presence of cholera toxin, ST and SNP typing of CC69, can all be derived from a single process work flow. WGS data may also be interrogated for additional virulence factors, and antimicrobial resistance determinants. The genomic data of all *V. cholerae* sequenced at PHE are publically released into the NCBI BioProject PRJNA438219 in order to facilitate public health surveillance, and monitoring of the global transmission of the pandemic lineages, by the international scientific community.

---

Supplementary Figure 1. Phylogeny of ST69 comprising isolates from this study (highlighted in red) and from publicly available databases

Supplementary Table 1. Short read archive accessions, WGS data and travel data for the sequenced isolates (n=152)

## References

1. Harris JB, LaRocque RC, Qadri F, Ryan ET, Calderwood SB. 2012. Cholera. Lancet 379:2466–76.

2. Jutla A, Khan R, Colwell R. 2017. Natural Disasters and Cholera Outbreaks: Current Understanding and Future Outlook. Curr Environ Health Rep 4:99–107.

3. Islam MT, Alam M, Boucher Y. 2017. Emergence, ecology and dispersal of the pandemic generating Vibrio cholerae lineage. Int Microbiol 20:106–115.

4. Crowe SJ, Newton AE, Gould LH, Parsons MB, Stroika S, Bopp CA, Freeman M, Greene K, Mahon BE. 2016. Vibriosis, not cholera: toxigenic Vibrio cholerae non-O1, non-O139 infections in the United States, 1984-2014. Epidemiol Infect 144:3335–3341.

5. Mutreja A, Kim DW, Thomson NR, Connor TR, Lee JH, Kariuki S, Croucher NJ, Choi SY, Harris SR, Lebens M, Niyogi SK, Kim EJ, Ramamurthy T, Chun J, Wood JL, Clemens JD, Czerkinsky C, Nair GB, Holmgren J, Parkhill J, Dougan G. 2011. Evidence for several waves of global transmission in the seventh cholera pandemic. Nature 477:462–5.

6. Albert MJ. 1996. Epidemiology & molecular biology of Vibrio cholerae O139 Bengal. Indian J Med Res 104:14–27.

7. Clemens JD, Nair GB, Ahmed T, Qadri F, Holmgren J. 2017. Cholera. Lancet 390:1539–1549.

8. Domman D, Quilici ML, Dorman MJ, Njamkepo E, Mutreja A, Mather AE, Delgado G, Morales-Espinosa R, Grimont PAD, Lizárraga-Partida ML, Bouchier C, Aanensen DM, Kuri-Morales P, Tarr CL, Dougan G, Parkhill J, Campos J, Cravioto A, Weill FX, Thomson NR. 2017. Integrated view of Vibrio cholerae in the Americas. Science 358:789–793.

9. Weill FX, Domman D, Njamkepo E, Tarr C, Rauzier J, Fawal N, Keddy KH, Salje H, Moore S, Mukhopadhyay AK, Bercion R, Luquero FJ, Ngandjio A, Dosso M, Monakhova E, Garin B, Bouchier C, Pazzani C, Mutreja A, Grunow R, Sidikou F, Bonte L, Breurec S, Damian M, Njanpop-Lafourcade BM, Sapriel G, Page AL, Hamze M, Henkens M, Chowdhury G, Mengel M, Koeck JL, Fournier JM, Dougan G, Grimont PAD, Parkhill J, Holt KE, Piarroux R, Ramamurthy T, Quilici ML, Thomson NR. 2017. Genomic history of the seventh pandemic of cholera in Africa. Science 358:785–789.

10. Dallman TJ, Byrne L, Ashton PM, Cowley LA, Perry NT, Adak G, Petrovska L, Ellis RJ, Elson R, Underwood A, Green J, Hanage WP, Jenkins C, Grant K, Wain J. 2015. Whole-genome sequencing for national surveillance of Shiga toxin-producing Escherichia coli O157. Clin Infect Dis 61:305–12.

11. Ashton PM, Nair S, Peters TM, Bale JA, Powell DG, Painset A, Tewolde R, Schaefer U, Jenkins C, Dallman TJ, de Pinna EM, Grant KA; Salmonella Whole Genome Sequencing Implementation Group. 2016. Identification of Salmonella for public health surveillance using whole genome sequencing. PeerJ 4:e1752.

12. Chattaway MA, Greig DR, Gentle A, Hartman HB, Dallman TJ, Jenkins C. 2017. Whole-Genome Sequencing for National Surveillance of Shigella flexneri. Front Microbiol 8:1700.

13. Bolger AM, Lohse M, Usadel B. 2014. Trimmomatic: A flexible trimmer for Illumina Sequence Data. Bioinformatics 30:2114–20.

14. Chattaway MA, Schaefer U, Tewolde R, Dallman TJ, Jenkins C. 2017. Identification of Escherichia coli and Shigella species from Whole-Genome Sequences. J Clin Microbiol 55:616–623.

15. Tewolde R, Dallman T, Schaefer U, Sheppard CL, Ashton P, Pichon B, Ellington M, Swift C, Green J, Underwood A. 2016. MOST: a modified MLST typing tool based on short read sequencing. PeerJ 4:e2308.

16. Li H, Durbin R. 2010. Fast and accurate long-read alignment with Burrows-Wheeler transform. Bioinformatics 26:589–595.

17. Bankevich A, Nurk S, Antipov D, Gurevich AA, Dvorkin M, Kulikov AS, Lesin VM, Nikolenko SI, Pham S, Prjibelski AD, Pyshkin AV, Sirotkin AV, Vyahhi N, Tesler G, Alekseyev MA, Pevzner PA. 2012. SPAdes: a new genome assembly algorithm and its applications to single-cell sequencing. J Comput Biol 19:455–477

18. McKenna A, Hanna M, Banks E, Sivachenko A, Cibulskis K, Kernytsky A, Garimella K, Altshuler D, Gabriel S, Daly M, DePristo MA. 2010 The Genome Analysis Toolkit: a MapReduce framework for analyzing next-generation DNA sequencing data. Genome Research 20:1297–1303.

19. Stamatakis A. 2014 RAxML version 8: a tool for phylogenetic analysis and postanalysis of large phylogenies. Bioinformatics 30:1312–1313.

20. Langmead B, Salzberg SL. 2012. Fast gapped-read alignment with Bowtie 2. Nat Methods 9:357–359.

21. Greig DR, Hickey TJ, Boxall MD, Begum H, Gentle A, Jenkins C, Chattaway MA. A real-time multiplex PCR for the identification and typing of Vibrio cholerae. Diagn Microbiol Infect Dis 90:171–176.

22. Mukhopadhyay AK, Takeda Y, Balakrish Nair G. 2014. Cholera outbreaks in the El Tor biotype era and the impact of the new El Tor variants. Curr Top Microbiol Immunol 379:17–47.

23. Siddique AK, Cash R. 2014. Cholera outbreaks in the classical biotype era. Curr Top Microbiol Immunol 379:1–16.

24. Ghosh R, Sharma NC, Halder K, Bhadra RK, Chowdhury G, Pazhani GP, Shinoda S, Mukhopadhyay AK, Nair GB, Ramamurthy T. 2016. Phenotypic and Genetic Heterogeneity in Vibrio cholerae O139 Isolated from Cholera Cases in Delhi, India during 2001-2006. Front Microbiol 7:1250.

25. Anandan S, Devanga Ragupathi NK, Muthuirulandi Sethuvel DP, Thangamani S, Veeraraghavan B. 2017. Prevailing clone (ST69) of Vibrio cholerae O139 in India over 10 years. Gut Pathog. 9:60.

26. Luo Y, Octavia S, Jin D, Ye J, Miao Z, Jiang T, Xia S, Lan R. 2016. US Gulf-like toxigenic O1 Vibrio cholerae causing sporadic cholera outbreaks in China. J Infect 72:564–72.

27. Ramamurthy T, Sharma NC. 2014. Cholera outbreaks in India. Curr Top Microbiol Immunol 379:49–85.

28. Shah MA, Mutreja A, Thomson N, Baker S, Parkhill J, Dougan G, Bokhari H, Wren BW. 2014. Genomic epidemiology of Vibrio cholerae O1 associated with floods, Pakistan, 2010. Emerg Infect Dis 20:13–20.

29. Kotetishvili M, Stine OC, Chen Y, Kreger A, Sulakvelidze A, Sozhamannan S, Morris JG Jr. 2003. Multilocus sequence typing has better discriminatory ability for typing Vibrio cholerae than does pulsed-field gel electrophoresis and provides a measure of phylogenetic relatedness. J Clin Microbiol 41:2191–6

30. Lam C, Octavia S, Reeves PR, Lan R. 2012. Multi-locus variable number tandem repeat analysis of 7th pandemic Vibrio cholerae. BMC Microbiol 12:82.

31. Octavia S, Salim A, Kurniawan J, Lam C, Leung Q, Ahsan S, Reeves PR, Nair GB, Lan R. 2013. Population structure and evolution of non-O1/non-O139 Vibrio cholerae by multilocus sequence typing. PLoS One 8:e65342.

32. Siriphap A, Leekitcharoenphon P, Kaas RS, Theethakaew C, Aarestrup FM, Sutheinkul O, Hendriksen RS. 2017. Characterization and genetic variation of Vibrio cholerae isolated from clinical and environmental sources in Thailand. PLoS One 12:e0169324.

33. Chowdhury FR, Nur Z, Hassan N, von Seidlein L, Dunachie S. 2017. Pandemics, pathogenicity and changing molecular epidemiology of cholera in the era of global warming. Ann Clin Microbiol Antimicrob 16:10.

34. Neilson AA, Mayer CA. 2010. Cholera - recommendations for prevention in travellers. Aust Fam Physician 39:220–6.

